# Vitamin D counters bone invasion by mammary cancer through inhibition of inflammation and epithelial-to-mesenchymal transition

**DOI:** 10.64898/2026.09.02.748895

**Authors:** Jiarong Li, Aimée-Lee Luco, Anne Camirand, Richard Kremer

## Abstract

Vitamin D deficiency is associated with poor outcome in several cancers in humans, and administration of vitamin D or analogs has been shown to decrease tumor progression and metastasis in animal mammary cancer models. We previously demonstrated significant acceleration of carcinogenesis in vitamin D-deficient mouse mammary tumor virus-polyoma middle T (MMTV-PyMT) mammary cancer model as well as of its spontaneous metastasis to lungs. While vitamin D also plays a role in skeletal metastasis, detailed mechanisms of its promotion of bone invasion and metastatic events are not completely elucidated. In the present study we used tibially-injected MMTV-PyMT mammary tumor cells to analyse how dietary-induced vitamin D deficiency in non-immunodeficient FVB mice accelerates bone invasion. Mechanistically, we observed vitamin D deficiency to increase pro-inflammation cytokines and nestin expression in internal bone surface and marrow, and to increase epithelial-to-mesenchymal transition (EMT) through Zeb1 transcription factor. *In vitro*, treatment of MMTV-PyMT tumor cells with CXCL12 was observed to stimulate Zeb1 expression, and this effect was efficiently countered by 1,25(OH)_2_D treatment. Analysis of cytokines in MMTV-PyMT mammary tumor cells *in vitro* showed significant reduction in several pro-inflammatory cytokines with 1,25(OH)_2_D treatment (GM-CSF, ICAM-1, IL-1ra, IP-10, JE, MCP-5, MIP-1α, MIP-1β, MIP-2, RANTES and CXCL12), a crucial observation in view of the current evidence that inflammation is one of the hallmarks of cancer. Furthermore, vitamin D repleteness is associated with very high expression of Socs1 (suppressor of cytokine signalling 1), an inhibitor of JAK/STAT pathway which prevents excessive inflammatory responses and has a tumor-suppressive role. These findings provide a strong link between vitamin D deficiency and acceleration of inflammation-driven bone invasion, and nestin and EMT. The evidence suggests that vitamin D-repleteness in breast cancer patients could enhance the efficacy of co-administered therapies in preventing invasion of skeletal sites.

**LAY SUMMARY:** Vitamin D deficiency, a very common occurrence in the modern world, is associated with poor outcome for several cancers among which colorectal, breast, kidney and thyroid. We previously showed that a vitamin D-deficient diet caused significant acceleration of carcinogenesis and lung metastasis in mouse. Vitamin D also plays a role in skeletal metastasis, however, mechanisms are not yet elucidated. Here, we demonstrate that vitamin D deficiency acts through pro-inflammation factors to help skeletal progression of mammary cancer. Evidence suggests that adequate vitamin D status in breast cancer patients could enhance efficacy of co-administered therapies in preventing deadly invasion of bone.

## INTRODUCTION

Vitamin D is a potent steroid hormone which affects differentiation, proliferation and apoptosis mechanisms in almost every cell in the body [1, 2]. Humans obtain some vitD in their diet but the main input occurs through synthesis in skin exposed to sunlight. In this process, the precursor 7-dihydrocholesterol is transformed by UVB rays into pre-vitD_3_ and cholecalciferol, and a series of hydroxylations produce the biologically-active vitD metabolite 1,25(OH)_2_D in the liver, kidneys and extra-renal tissues. 1,25(OH)_2_ D binds the vitD receptor (VDR) which heterodimerizes with the retinoid X receptor, allowing interaction with vitamin D-responsive DNA elements and regulation of expression of hundreds of genes among which those involved in bone mineralization, calcium and phosphorus homeostasis [3]. The vitD axis also regulates genes involved in immune response, including those inhibiting the production of pro-inflammatory cytokines and promoting the synthesis of anti-inflammatory cytokines [4].

Compared to these well-known roles for vitD, a non-classical action in cancer is a more recent but crucial observation [5]. VitD displays anti-proliferative effects in many animal cancer models [6], and many studies have shown a direct association between low precursor 25(OH)D in serum and a risk for colon, breast, prostate, and gastric cancers, among others [7], as well as an increased risk of mortality [6–16]. We showed chemoprevention activity of precursor 25(OH)D in the PyMT MMTV mouse model of mammary cancer [17] and the crucial role of tumor-produced 1,25(OH)_2_D in tumor progression in the same model [18]. The antitumor activity of vitD is associated with its ability to inhibit cell cycle progression [19], tumor growth [18, 20, 21], angiogenesis [22], apoptosis [23, 24] and epithelial to mesenchymal transition (EMT)[7, 25–27].

It is important to remember that most patients do not die from the primary tumor but because of cancerous invasion to distal sites. Metastatic expansion of cancer involves migration of primary tumor cells to distal sites where invasion results in deadly consequences. A frequent metastatic site for cancer metastasis is bone, which is presently an impossible target to cure. In immunodeficient rodent models, vitD deficiency is associated with accelerated bone metastasis [28–30], and in humans, a high prevalence of vitD deficiency has been reported in patients with metastatic bone disease derived from multiple myeloma, the most common cancer to produce bone lesions, as well as from breast and prostate cancer [31]. Our previous work illustrated the important effects of in-tumor synthesis of 1,25(OH)_2_D against tumor initiation, growth and metastasis in the MMTV-PyMT mouse model of mammary cancer [32]. With respect to following metastatic events, we demonstrated that vitD regulates CXCL12/CXCR4 interactions and EMT in lung metastasis of mammary tumor cells in the MMTV-PyMT mouse model [33]. We also demonstrated the efficacy of the low calcemic vitD analog EB1089 in preventing bone metastasis by MDA-MB-231 human breast cancer cells injected in the heart of nude mice, and in prolonging the animals survival time [34]. Dietary vitD deficiency has been observed to facilitate bone invasion by human MDA-MB-231 breast cancer cells in a mouse model of malignant skeletal metastasis [28–30], and metastasis to the liver was enhanced in mice injected in the mammary fat pad with VDR-ablated mammary tumor cells [35]. Growing evidence therefore points to a clear involvement of vitD in metastasis, but by mechanisms that are not completely elucidated. We show in the present study that these mechanisms include the action of inflammatory cytokines, nestin, chemokines, and EMT-promoting Zeb1 transcription factor.

Nestin (neuroepithelial stem cell protein) filament protein is found in a broad variety of cells such as proliferating vascular endothelial cells, basal cells of the mammary glands, and liver stem cells [36–38]. It is also highly-expressed in many high-metastatic cancers including breast, and is associated with aggressiveness and poor prognosis. Nestin mechanism of action in tumor progression and metastasis involves its role in reducing the cell stiffness controlling tumor-associated angiogenesis [39, 40]. With high nestin expression, new blood vessels display fragile basement membranes, thus allowing tumor cells to enter the circulation and disseminate [41]. Nestin knock-out in tumor cells decreases metastasis [42] and is consequently speculated as a new anti-metastatic therapeutic strategy.

Chemokines are a family of small proteins (8 -10 kDa) with chemoattractant properties that direct the migration of cells presenting the appropriate G-protein coupled receptor on their surface. CXCL12 (also known as stromal cell-derived factor-1 or SDF1) is a universally-expressed chemokine involved in embryogenesis, angiogenesis and inflammation events, in pathological states such as HIV infection, and is particularly enriched in metastasis target sites such as bone, lung and liver. It is also involved in progression of many cancers including breast [43]. CXCL12 as a ligand binds the CXCR4 receptor present on tumor cells, which contributes to tumor cell immobilisation, growth, angiogenesis, therapeutic resistance and metastasis [44], therefore CXCL12 expression correlates with poor prognosis in human breast cancer patients [45–47]. Proinflammatory cytokines that modulate the immune responses also cause tumor progression and promote invasion and metastasis through a great variety of mechanisms [48].

Zeb1 (Zinc finger E-box binding homeobox 1) transcription factor drives cancer metastasis by inducing epithelial-to-mesenchymal transition (EMT) in tumor cells. EMT is a cellular mechanism involved in normal physiological states and in pathological conditions such as wound healing, tissue fibrosis, and cancer [49]. Zeb1 represses epithelial genes such as E-cadherin, causing cells to lose adhesion and gain migratory mesenchymal properties, therefore promoting invasion, migration, and resistance to therapy in numerous cancers [50]. High Zeb1 levels promote mammary epithelial tumors [51] and often correlate with increased metastatic risk and poor prognosis [52]. Targeting Zeb1 or its associated signaling pathways can reduce metastasis, reverse chemoresistance, and restore sensitivity to therapies [53].

Once breast cancer metastases are established in target organs such as bone or lung, the condition is generally considered incurable. There is therefore an urgent need to improve current therapies that address cancer spread. The present study aims to investigate mechanistic aspects of bone tumoral invasion acceleration by vitD deficiency. To allow observation of the role of components from the immune system, we used non-immunodeficient mice that received allografts of mammary tumor cells from the MMTV-PyMT mouse model of mammary carcinogenesis, and we compared bone invasion in vitD-deficient and vitD-replete non-immunodeficient FVB animals.

## MATERIALS AND METHODS

### Animal breeding and diet

PyMT-MMTV *Cyp27b1^flox/flox^*MMTV-Cre and *Cyp27b1^wt/wt^* MMTV-Cre mice were produced as described previously [18]. Non-immunodeficient FVB mice (female) were purchased from Charles River Laboratories (Senneville, QC). Three weeks after birth, pups were weaned and then fed *ad libitum* a vitD-deficient diet (25 IU/Kg) or a normal vitD diet (1000 IU/Kg)(Harlan, Montréal, QC). Mice were kept in a room without UVB lighting in order to minimize the endogenous production of vitD in the skin.

### PyMT MMTV mammary tumor cell isolation, culture and analysis

Mammary tumors were dissected, minced and digested in DMEM without fetal bovine serum (FBS) containing 2.4 mg/ml collagenase B and dispase II (Roche Canada, QC). Floating cells were washed, pelleted, resuspended and filtered through a 40 µm sterile nylon mesh strainer then plated in pre-warmed DMEM medium with 10% FBS and cultured in 6-well or 10 cm plates at initial density of 3 × 10^6^ cells/ml. After the first passage, cells at 100% confluency were trypsinized, aliquoted and stored in liquid nitrogen.

For 1, 25(OH)_2_D and CXCL12 treatment experiments, cells were seeded in 6-well plates and cultured at 37^0^C, 5% CO_2_, and 95% humidity in DMEM containing 10% FBS. At 70% confluency, cells were starved overnight in DMEM without FBS, treated with 10^−7^ M 1,25(OH)_2_D (Sigma), CXCL12 (100ng/ml, BIO-RAD Canada) or vector (ethanol) in 7% charcoal-stripped FBS (Invitrogen, ON). The treated cells were either used for immunofluorescence staining and confocal microscopy or harvested for RNA analysis. The experiments were conducted in duplicates.

### Intratibial injection of mouse MMTV PyMT mammary tumor cells

Mouse MMTV-PyMT mammary tumor cells were cultured to 100% confluency in DMEM containing 10% FBS, trypsinized and resuspended in PBS at a concentration of 5×10^5^ cells/15µl. Intratibial injections were performed on 8-week-old FVB female mice as follows: the proximal end of the left tibia was exposed surgically and the knee was maintained in a flexed position. While grasping the ankle/leg of the mouse, a 25G needle was inserted as a guide and a 30G needle inserted inside the 25G to inject a 15 µl cell suspension. The animals were sacrificed five weeks later (age 13 weeks) and bones were fixed as described below.

### Detection of bone lesions by X-ray

At sacrifice, animals were placed in the supine position in an imaging tray and X-rays of the hind limbs were taken with a Bruker In-Vivo Xtreme optical imager (Center for Bone and Periodontal Research, McGill University). X-rays were analyzed for the presence of bony lesions and scored according to the following method: 0 (no lesion), 1 (minor changes), 2 (small lesions), 3 (significant lesions with minor peripheral margin breaks) and 4 (significant lesions with major peripheral margin breaks and > 10% of bone surface disrupted). The X-rays were independently scored by two individuals who did not know which group the mice belonged to.

### Micro-computed tomography (Micro CT)

Tibias were dissected free of soft tissue, fixed overnight in 70% ethanol and analyzed by micro-CT with a SkyScan 1072 scanner (Center for Bone and Periodontal Research, McGill University) and associated analysis software (SkyScan, Antwerp, Belgium).

### Histology and confocal microscopy

Immunofluorescence staining was conducted according to the manufacturer’s instructions (Invitrogen, ON). Briefly, after antigen unmasking, the specimens were incubated in 10% serum in PBS for 30 minutes, washed 3 min in PBS, and incubated overnight at 4°C with rabbit anti-CXCL12 antibody (PA5-89116, Invitrogen Burlington, ON) or anti-mouse CXCR4 mAb (MAB21651, R&D). The slides were washed 5 minutes with PBST and incubated 45 minutes in a dark chamber with the fluorochrome-conjugated secondary antibody (donkey anti-rabbit conjugated with Alexa fluor 488 (A-11008) and goat anti-mouse conjugated Alexa fluor 568 (A-11004) (Invitrogen, Burlington, ON). Slides were washed and counterstained 10 minutes in the dark with 4’,6-diamino-2-phenylindole (DAPI)(Invitrogen, ON), washed with three changes of PBS and mounted under coverslips in aqueous mounting medium (Thermo Electron, Pittsburgh, PA). Results were analyzed with an LSM 880 Meta confocal microscope (Carl Zeiss MicroImaging Gmbh, Munich, Germany). Some bones were fixed in 70% ethanol overnight at 4°C, rinsed in PBS and decalcified in EDTA glycerol solution for 7-10 days at 4 °C. Decalcified bones were dehydrated and paraffin-embedded, and 5-μm sections were stained for immunofluorescence (IF) or with H & E for CXCL12 and CXCR4 as described above.

### Laser scanning confocal and super-resolution microscopy

A Zeiss LSM 880 with ElyraPS1 laser scanning confocal and super-resolution microscope system was used (Carl Zeiss, Jena, Germany; MUHC, RI molecular imaging platform). The ElyraPS1 laser scanning image microscope acquires a series of images producing a “Z stack”. The raw and processed images (confocal and ElyraPS1 laser scanning) were analyzed by the Zeiss Zen software. Maximum intensity projections and orthogonal sections were produced for each image using the Zen software package. Staining for CXCR4 and CXCL12 was done as above.

### RT^2^ Profiler PCR arrays assay

Total RNA was extracted from tissues using Trizol reagent (Thermo Fisher Scientific, Canada) according to the manufacturer’s instructions. RNA quality was measured by spectrophotometer, and RNA quality control parameters OD260/280 were between 1.8–2.0. Reverse transcription was performed using the All-in-One™ First-Strand cDNA Synthesis Kit (5X all-in-one RT masterMix Cat G490) following the manufacturer’s procedure. RT2 Profiler PCR arrays plate (QIAGEN, catalog number PAMM-039Y) in combination with RT2 SYBR® Green qPCR Mastermix (catalog number 330,503) were performed for cDNA real time PCR. Each array plate contained a set of 96 wells for testing. Genomic DNA contamination, reverse transcription, and positive PCR controls were included in each 96-well set on each plate. Glyceraldehyde-3-phosphate dehydrogenase (GAPDH) was used as a reference gene for detection. CT values were exported to an Excel file to construct a CT value table, and uploaded to data analysis portal http://www.qiagen.com/geneglobe. Samples included control and test groups. CT values were normalized based on automatic selection from a full set of reference genes.

### Proteome profiler mouse cytokine array assay

PYMT mouse mammary cell culture at 70% confluency were starved overnight in DMEM without FBS, treated with 10^−7^ M 1,25 (OH)_2_D (Sigma) or vector (ethanol) in 7% charcoal-stripped FBS (Invitrogen, ON). Supernatants were collected from tumor cell cultures after 24 hour treatment. The supernatants were filtered through 0.2mm syringe filters (Corning) and the resulting tumor-conditioned media were stored in aliquots at 80°C. The Proteome Profiler Mouse Cytokine Array Kit (ARY006, Panel A; R&D Systems, Minneapolis, MN, USA) was used to quantify 40 different cyto- and chemokines in the supernatants, according to the manufacturer’s instructions (**supplementary figure 1**). Digitalized radiographic films were analyzed using QuickSpots densitometer software (R&D System) and values expressed as relative pixel density.

### RNA analysis

RNA was extracted with a Qiagen kit (Qiagen, Canada). The cDNA RT kit and SYBR green master mix were from Roche Canada (Laval, QC). The 2^−Δ Δ Ct^ method was used to analyze the results. Forward and reverse primers for amplification of mouse mRNAs were:

CXCL12: 5′-CAGAGCCAACGTCAAGCA-3’ and 5’-CTTGTCTACCTCTACCCCGACAT-3’

Zeb1: 5’-TGCACTGAGTGTGGAAAAGC-3’ and 5’-TGGTGATGCTGAAAGAGACG-3’

GAPDH: 5’-AACGACCCCTTCATTGAC-3’ and 5’-TCCACGACATACTCAGCAC-3’.

### Statistics

The statistical difference of tumor onset rate of the animals was determined by Kaplan-Meier analysis. For tumor progression, numerical data are presented as mean ± SD. Data were analyzed by ANOVA followed by a Bonferri *post hoc* test to determine the statistical significance of the differences. All statistical analyses were performed using Graph Pad Software and P < 0.05 was considered statistically significant. For qRT-PCR data, statistical analyses were performed (n =3). This package uses ΔΔCT based fold change calculations and the Student’s t test to calculate 2-tail, equal variance P values. P <0.05 and a fold change greater than 1.5 were considered to be a significant difference.

### Animal Ethics

All animal studies were carried out in compliance with regulations of the McGill University institutional animal care committee. All animal surgeries were conducted in accordance with principles and procedures dictated by the highest standards of humane animal care.

## RESULTS

### Vitamin D dietary deficiency promotes mammary tumor cell growth in bone *in vivo*

Female FVB mice were fed a normal (1000 IU/kg) or low (25 IU/kg) vitD diet from weaning. At 8 weeks, 5×10^5^ MMTV-PyMT mammary tumors cells were injected intratibially. At sacrifice (6 weeks later) bone histomorphometry revealed a striking difference in skeletal tumoral invasion between vitD deficient and vitD-replete animals (**Figure 1 a,b,c**). The average surface of bone lesions was 158.4 ± 35.1mm^2^ for mice on the 1000 IU diet but 596 ± 113 mm^2^for animals on the 25 IU diet. In micro-CT analysis, vitD-replete animals displayed (BV/TV) 9.68 ± 1.42% bone volume loss compared to 28.2 ± 2.3% for deficient mice (**Figure 1 e, f**). These results suggest that sufficient levels of dietary vitD can significantly reduce the progression of skeletal invasion by allografted mammary tumor cells.

**Figure 1:**
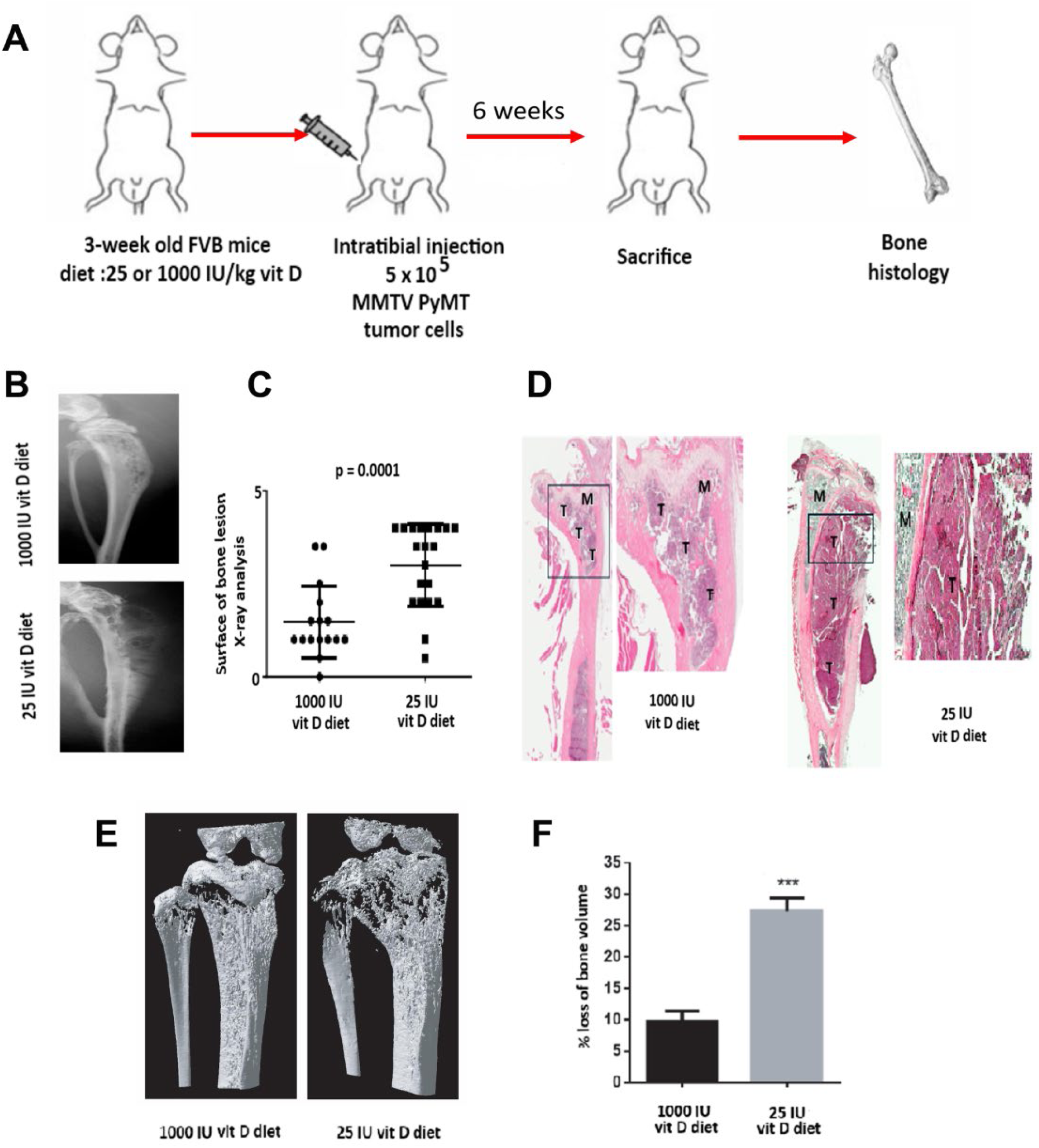
Vitamin D dietary deficiency promotes mammary tumor cell growth in bone *in vivo*. (**A**) Experimental design: at weaning (3-week old) female FVB mice were fed a normal (1000 IU/kg) or low (25 IU/kg) vitamin D3 diet. At 8 weeks, 5×10^5^ MMTV-PyMT mammary tumors cells were injected intratibially. Mice were sacrificed 6 weeks later and bone imaging histomorphometry performed. (**B**) Tibial X-ray scans. T: tumor, M: marrow. (**C**) Scoring of bone lesions on X-rays. (**D**) Histology (H and E) of decalcified paraffin-embedded tibial bone sections of mice on normal and low vitD diet. Tumoral area in tibiae of mice on normal or low vitD diet. (**E**) Representative micro-CT images of bone. (**F**) Bone volume loss by micro-CT at sacrifice. There were 37 animals in the normal vitD group and 41 animals in the low vitD group. p <0.001 indicates a significant difference between a low vs normal vitamin D diet. This experiment was repeated twice and results were combined.

### Vitamin D dietary deficiency increases nestin expression on internal surface of bones and in flushed bone marrow of tumor cell-injected bones

Immunofluorescence and real time PCR analysis indicate that dietary deficiency in vitD significantly increases skeletal relative expression of nestin mRNA levels fold changes in vitamin D replete animals are 1.98 ± 0.03 vs 1.0 ± 0.016 in vitD deficient mice (**Figure 2 a,b**). These results indicate a significant difference between levels of expression of nestin on internal surface of bones and in flushed bone marrow of tumor cell-injected bones in normal and low vitD diet group.

**Figure 2:**
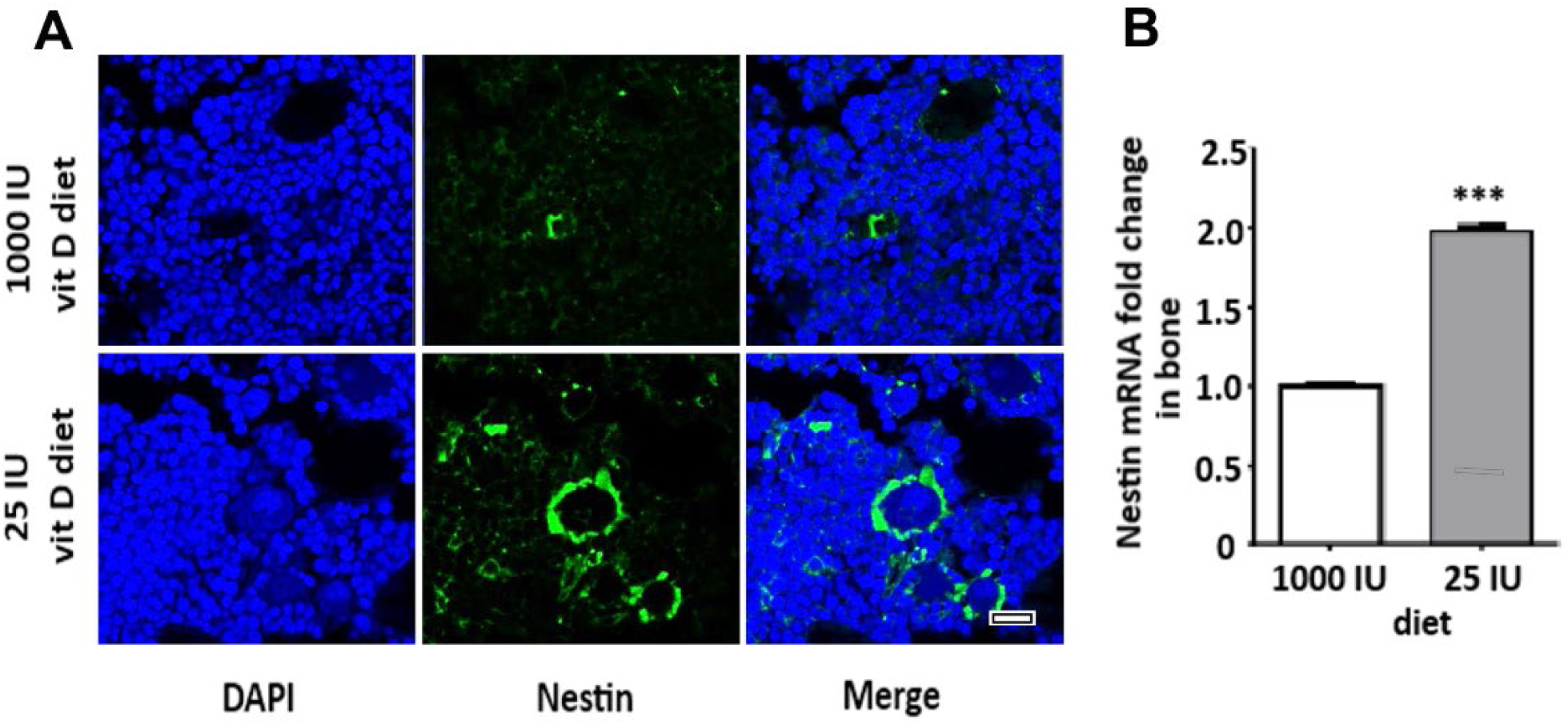
Vitamin D dietary deficiency increases nestin expression on internal bone surface and in flushed bone marrow. **(A)** IF detection of nestin expression on internal bone surface of mice on normal (1000 IU) or low (25 IU) vitD diet. (**B)** Real-time PCR of nestin mRNA in flushed bone marrow of mice on normal or low vitD diet. *** P <0.001 indicates a significant difference between normal and low vitD diet group. Scale bar =100μm

### Vitamin D dietary deficiency increases CXCL12 and CXCR4 expression and their co-localisation on internal bone surface and flushed bone marrow

Immunofluorescence analysis and real time PCR indicate that dietary deficiency in vitD significantly increases relative expression of CXCL12 mRNA levels fold changes (**Figure 3 a,b**) and CXCR4 (**Figure 3 c,d**). mRNA levels for CXCL12 in vitD replete animals are 1.89 ± 0.08 vs 1.0 ± 0.072 in vitD mice. mRNA levels for CXCR4 in vitD replete animals are 2.68 ± 0.26 and 1.0 ± 0.11 in vitD mice. These results indicate a significant difference between levels of expression of CXCL12 and CXCR4 on internal bone surface (**figure 3 a,c**) and in flushed bone marrow (**figure 3 b,d**) of normal and low vitD diet groups of mice. CXCR4-CXCL12 co-localisation is also enhanced by vitD status (**figure 3e**).

**Figure 3.**
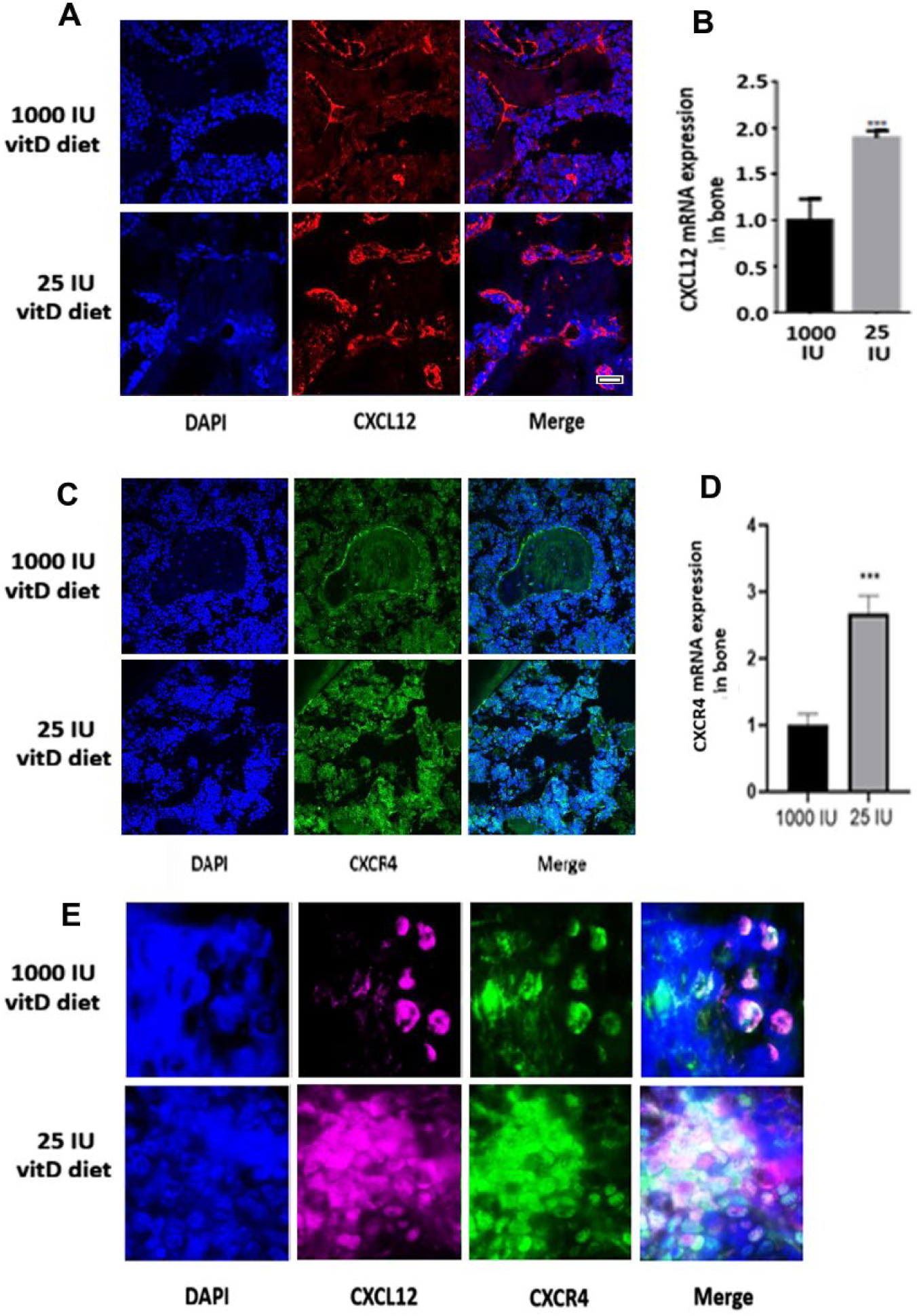
Vitamin D dietary deficiency increases CXCL12 and CXCR4 *in vivo,* on internal bone surface and flushed bone marrow. **(A)** IF detection of CXCL12 protein on internal surface of bone of mice on normal or low vitD diet. (**B**) Real-time PCR of CXCL12 mRNA in flushed bone marrow. (**C**) IF detection of CXCR4 protein on internal surface of bone of mice on normal or low vitD diet. (**D**) Real-time PCR of CXCR4 mRNA in flushed bone marrow (E) Co-localisation of CXCL12 and CXCR4 in flushed bone marrow. There were 32 animals in the 1000 IU/kg group and 36 animals in 25 the IU/kg group. *** P <0.001 indicates a significant difference between normal and low vitamin D3 diet group. scale bar =100μm.

### Vitamin D dietary deficiency increases bone Zeb1 and affects EMT markers vimentin and cadherin on internal surface of tumor-injected bone and flushed bone marrow *in vivo*

Immunofluorescence and real time PCR detection of Zeb1 protein in mice on normal or low vitD diet reveals a very important increase in expression of the transcription factor Zeb1 on the internal surface of bones and in flushed bone marrow (**Figure 4 a,b**). Vimentin and E-cadherin relative expression mRNA levels for vimentin in vitamin D replete animals are 35.8 ± 0.17 vs 1.0 ± 0.13 in vitamin D mice. mRNA levels for E-cadherin in vitD replete animals are 1.0 ± 0.12 and 2.36 ± 0.15 in vitamin D mice. These results indicate a significant difference in the EMT process between normal and low vitamin D3 diet groups where absence of sufficient dietary vitD causes a very significant augmentation of pro-EMT indicators.

**Figure 4.**
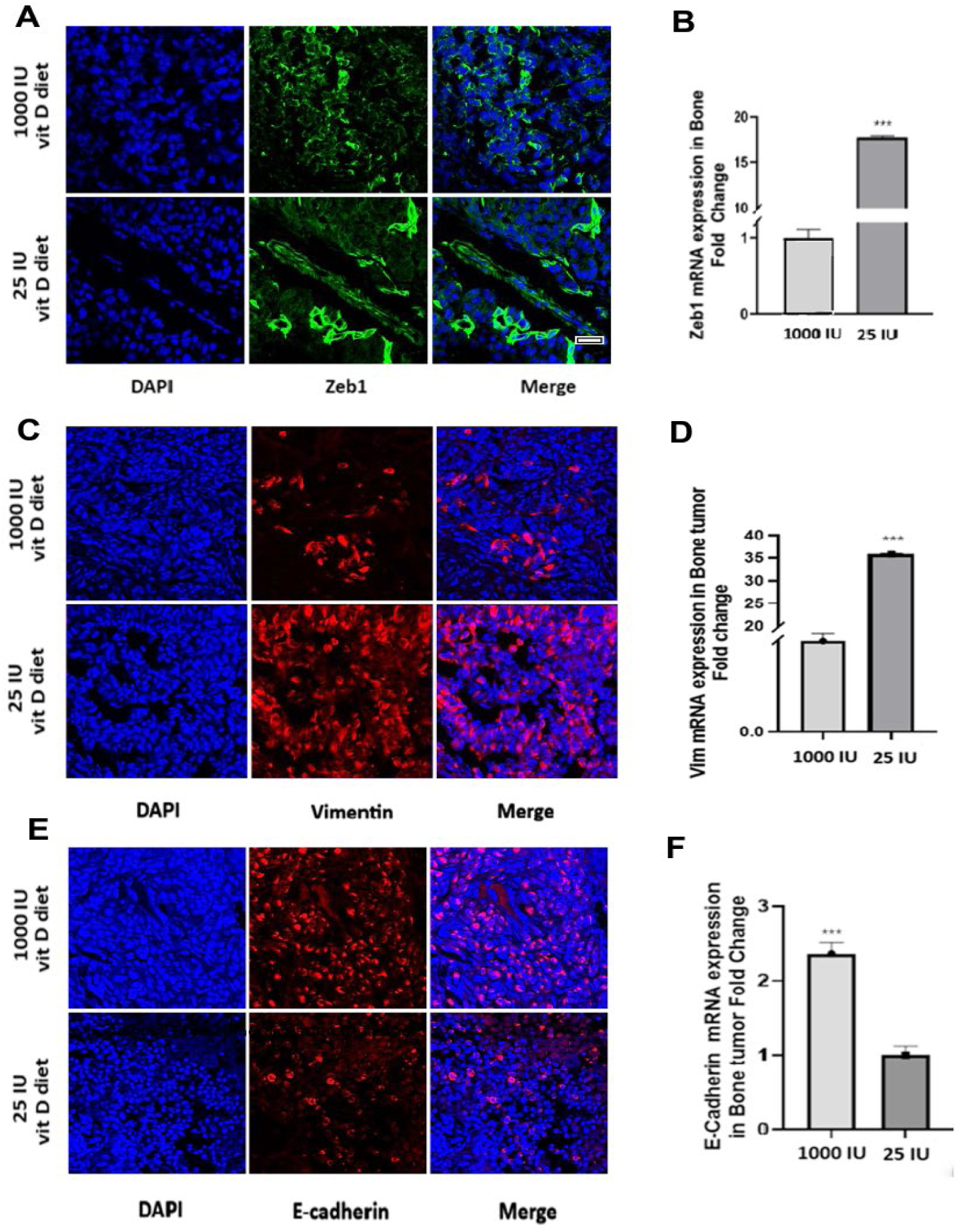
Vitamin D dietary deficiency increases bone Zeb1 and affects EMT markers on internal surface of tumor cell-injected bone and flushed bone marrow *in vivo.* (A) IF detection of Zeb1 protein on surface of bone of mice on normal or low vitD diet. (**B**) Real-time PCR of Zeb1 mRNA in flushed bone marrow. (C) Vimentin on surface of bone of mice on normal or low vitD diet. (D) real time PCR of vimentin mRNA (E) E-cadherin on surface of bone of mice on normal or low vitD diet. (F) Real time PCR of E-cadherin mRNA. There were 32 animals in the 1000 IU/kg group and 36 animals in 25 the IU/kg group. *** P <0.001 indicates a significant difference between normal and low vitamin D3 diet group. scale bar =100μm.

### 1, 25(OH)_2_D counters CXCL12 stimulation of EMT-deriving Zeb1 expression in MMTV PyMT mammary cancer cells *in vitro*

MMTV PyMT mammary tumor cells in culture were treated with either 1, 25(OH)_2_D (10^− 7^M) or CXCL12 (100ng/ml), or both. Cells were collected after 48 hours and immunofluorescence analysis as well as RT-PCT revealed a significant increase in expression of EMT-driving Zeb1 transcription factor due to CXCL12 mRNA fold change 3586 ± 153. This addition was very efficiently countered by addition of 1, 25(OH)_2_D (**Figure 5 a,b**). These results indicate that tumoral cells themselves boost their EMT-inducing Zeb1 levels when exogenous CXCL12 is present, and that 1, 25(OH)_2_D is able to inhibit this action.

**Figure 5:**
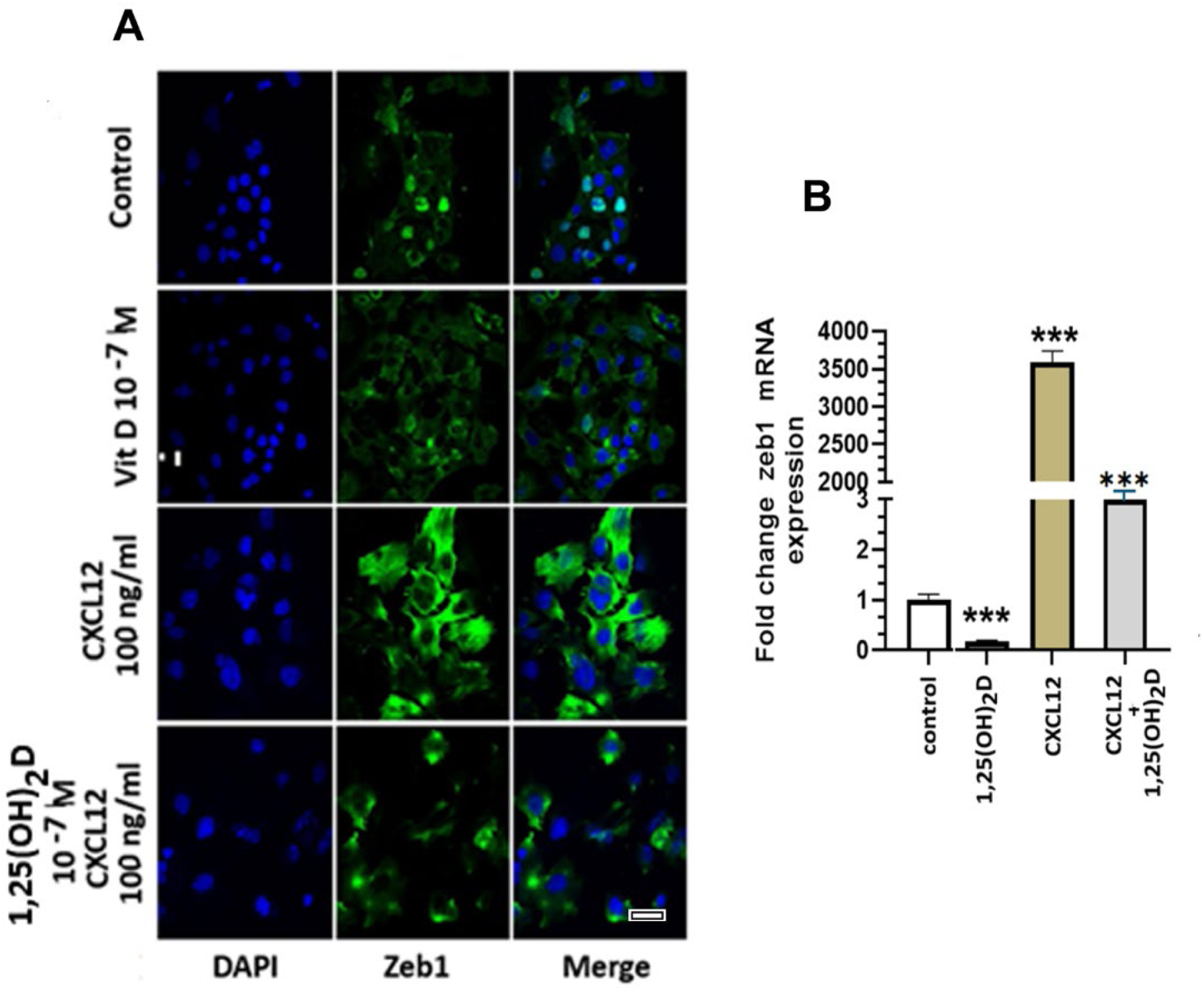
1,25 (OH)_2_D counters CXCL12 stimulation of Zeb1 expression in MMTV PyMT mammary cancer cells *in vitro:* **(A)** Zeb1 expression detected by confocal microscopy in MMTV PyMT tumor cells treated with vehicle or 1, 25(OH)_2_D (10^−7^M), CXCL12 (100ng/ml) or both. (**B**) Zeb1 mRNA expression in cells treated as above. Results are expression as fold change from vehicle vs treated cells. *** P <0.001 indicates a significant result. Scale bar =100um

### 1, 25(OH)_2_D inhibits MMTV-PyMT mouse mammary inflammatory cytokines *in vitro*

The Proteome Profiler Mouse Cytokine Array Kit was used to assess cell cytokine and chemokine profiles in tumor cell-conditioned supernatants. MMTV PyMT mammary tumor cells at 70% confluency in culture were starved overnight in DMEM without FBS, then treated with 10^− 7^ M 1,25 (OH)_2_D or vector (ethanol) in 7% charcoal-stripped FBS for 24 hours. Cell-conditioned supernatant was collected and analysed on the mRNA array for cytokines and chemokines (**Figure 6a**). For expression differences between 1, 25(OH)_2_D-induced mRNAs and vehicle-treated, boxes highlight genes showing statistical differences in expression between vehicle and 1, 25(OH)_2_D-treated (**Figure 6b**). 11 down-regulated genes were identified due 1,25(OH)_2_D treatment. Dots exhibiting stronger response density in vehicle compared to 1,25(OH)_2_D treatment are boxed in red. Histograms of pixel intensity for array values of boxed genes show that GM-CSF, ICAM-1, IL-1ra, IP-10, JE, MCP-5, MIP-1alpha, MIP-1beta, MIP-2, RANTES and CXCL12 spots remarkably decreased after 1,25(OH)_2_D 24 h treatment (**Figure 6c**). (A complete list of genes in the array is found in **supplementary figure S1**). Statistical difference was set as log2 fold change >2. These results indicate a strong ability of 1, 25(OH)_2_D to inhibit the expression of several pro-inflammatory cytokines, linking vitamin D anti-cancer activity to its inflammation control role.

**Figure 6:**
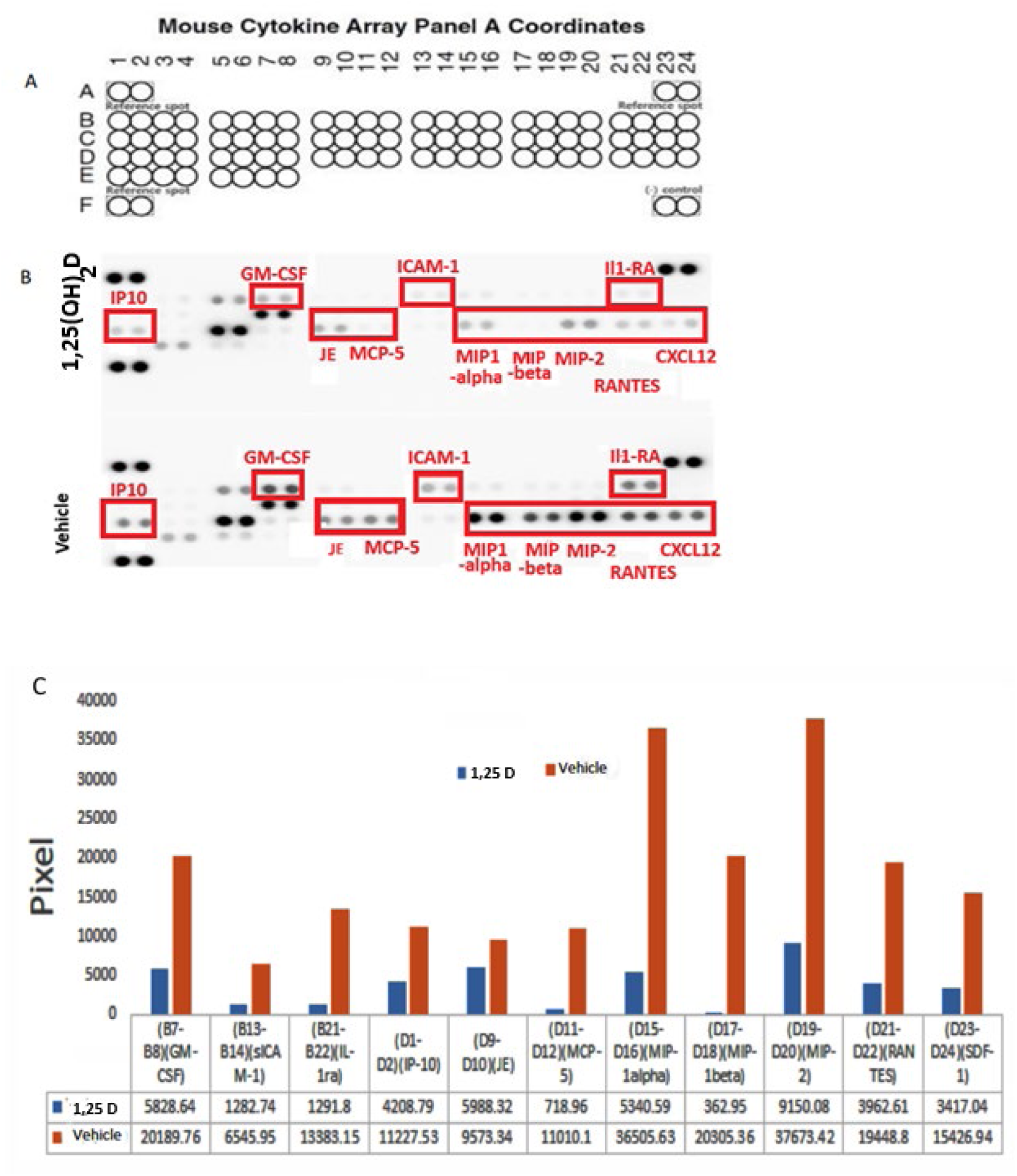
1, 25(OH)_2_D inhibits expression of MMTV-PyMT mouse mammary inflammatory cytokines in conditioned supernatant *in vitro.* Mouse tumor cell cytokine and chemokine profiles were assessed in conditioned supernatant using a Proteome Profiler Mouse Cytokine Array Kit kit after *in vitro* treatment of MMTV PyMT tumor cells with 1, 25(OH)_2_D (10^−7^ M) or vehicle. (**A**). Array outline. (Complete list in **supplementary figure S1**). (**B**) Protein expression difference between 1, 25(OH)_2_D -induced mRNAs and vehicle-treated cells. Boxes highlight cytokines showing statistical differences in expression between vehicle and 1, 25(OH)_2_D -treated. Statistical difference was set as log2 fold change >2. (**C**) Histograms of pixel intensity for array values of boxed genes show that GM-CSF, ICAM-1, IL-1ra, IP-10, JE, MCP-5, MIP-1alpha, MIP-1beta, MIP-2, RANTES and CXCL12 spots are remarkably decreased after 1,25(OH)_2_D 24 h treatment (blue: vit D treated, red: controls). Representative blots from 3 separate experiments.

### Vitamin D repleteness maintains MMTV PyMT mouse immune response signaling pathway

mRNA analysis on RT2 Profiler PCR array plate (**table 1**) indicates that vitD repleteness is associated with very high expression of Socs1 (suppressor of cytokine signalling 1). Socs1 is a negative regulator of cytokine signaling and inhibits the interferon-gamma (IFNγ) and JAK/STAT pathways. Along with Socs3 (also upregulated), it is involved in preventing excessive immune reactions. Among the other upregulated genes involved in regulation if the immune system and inflammation: Fcgr1 and A2m. Also upregulated by vitD repleteness are apoptosis controls Fas, Bcl2l1 and Smad3 and 4. In contrast, among genes downregulated by vitD repleteness are Nos2, Il2ra and Prl which are involved in inflammation and immune response. Oncogenes and proto-oncogenes Crk and Myc are also downregulated, as are genes involved in apoptosis Akt1 and cell mobility and invasion Csf1r. Taken together, these results suggest an implication of dietary vitD in the regulation of inflammation and immune pathways in cancer invasion of bone by mammary tumor cells.

**table 1:** Vitamin D repleteness maintains MMTV PyMT mouse immune response signaling pathway in flushed bone marrow. RT2 Profiler PCR arrays plate. Effect of vitD dietary deficiency compared to normal diet group mRNAs of signaling pathways. Only compounds with p < 0.05 difference are listed. Upregulated genes at top, downregulated genes below.

|  | Symbol | Description | fold change | p value | biological implication |
| --- | --- | --- | --- | --- | --- |
| UP | A2m | Alpha 2 macroglobulin | 6.82 | 0.004252 | immune response |
|  | Bcl2l1 | Bcl2-like 1 | 6.24 | 0.013692 | apoptosis |
|  | Fas | TNF receptor superfamily member 6 | 2.65 | 0.014845 | apoptosis |
|  | Fcgr1 | Fc receptor IgG, high affinity 1 | 5.25 | 0.008257 | immune response |
|  | Pias1 | Protein inhibitor of activated Stat1 | 8.5 | 0.007548 | JAK-STAT |
|  | Smad3 | MAD homolog 3 (Drosophila) | 3.62 | 0.035814 | tumor suppressor |
|  | Smad4 | MAD homolog 4 (Drosophila) | 4.22 | 0.024378 | tumor suppressor |
|  | Socs1 | Suppressor of cytokine signalling 1 | 170.58 | 0.003533 | immune response |
|  | Socs3 | Suppressor of cytokine signalling 3 | 10.28 | 0.012414 | immune response |
|  | Stam | Signal transducing adaptor molecule (SH3 domain and ITAM motif)1 | 5.12 | 0.022804 | JAK-STAT |
| DOWN | Akt1 | Thymoma viral protooncogene 1 | -1.56 | 0.04191 | apoptosis |
|  | Crk | V-Crk sarcoma virus CT10 oncogene homolog (avian) | -4.77 | 0.020319 | oncogene |
|  | Csf1r | Colony stimulating factor 1 receptor | -6.22 | 0.023773 | immune response |
|  | Il2ra | Interleukin 2 receptor alpha chain | -4.9 | 0.046723 | immune response |
|  | Myc | Myelocytomatosis oncogene | -5.93 | 0.022344 | oncogene |
|  | Nos2 | Nitric oxide synthase 2, inducible | -5.46 | 0.002573 | immune response |
|  | Prl | Prolactin | -5.09 | 0.015469 | immune response |

## DISCUSSION

Among the hallmarks of cancer, the phenomenon of inflammation has gained recognition as a critical component of the tumor microenvironment [48]. Like wounded tissues, tumors present high levels of cytokines, chemokines and their receptors, and have been described as ‘wounds that do not heal’, where continuous cell proliferation is driven by inflammation [54]. It is now known that inflammation promotes malignancy through the release of reactive oxygen species, proteases, cytokines and pro-angiogenic factors that cause vascularization, invasion and metastasis [55]. Because the skeletal environment is rich in components of the inflammatory pathways, it is an excellent target site for invasion by migrating tumor cells.

Vitamin D is a potent immune system modulator, actively suppressing the production of pro-inflammatory cytokines and rebalancing immune cell activity. Clinical studies suggest that serum 25(OH)D_3_ levels of less than 20 ng/mL (50 nmol/L) indicate vitD deficiency. Serum 25(OH)D_3_ levels below 30 ng/mL indicate insufficiency, while levels between 30 and 60 ng/mL (75 and 150 nmol/L) represent normal values. Epidemiological studies suggest an inverse association between circulating levels of 25(OH)D_3_ and inflammatory markers [56]. Maintaining optimal levels of vitD may therefore reduce chronic systemic inflammation and emerging evidence indicates that vitD deficiency plays a role in several inflammatory diseases among which: acute infections, cardiovascular disease, asthma, inflammatory bowel, chronic kidney and liver inflammatory disease, multiple sclerosis as well as inflammation/immune-related chronic disorders such as hypertension, diabetes, chronic lower-back pain, congestive heart failure, rheumatoid arthritis and systemic lupus erythematosus [56]. Observational studies suggest that vitD supplementation decreases biomarkers of inflammation in colorectal adenoma [57]and may protect against several infectious and auto-immune conditions [58]. It is known that 15-20% of all human tumors originate as a result of chronic infections and inflammatory conditions [59], however, the mechanisms by which vitD acts on the subsequent metastatic process are still unclear and may or may not involve inflammation. Previous work on vitD effect in bone metastasis involved xenografts in nude mice [28–30], a condition which does not allow the full immune response present in non-immunodeficient mice used in the present study. We show here the consequence of vitD repleteness in preventing the expression of several pro-inflammatory cytokines and chemokines: GM-CSF, ICAM-1, IL-1ra, IP-10, JE, MCP-5, MIP-1alpha, MIP-1beta, MIP-2, RANTES and CXCL12. We also observed stimulation of Socs1 and 3 and Plas1 expression, factors that prevent immune overreaction as known inhibitors of the JAK/STAT (Janus kinase-signal transducer and transcription) activator pathway which regulates physiological and pathological processes such as inflammation and stress [60, 61]. VitD repleteness also inhibits expression of the Crk (CT10 regulator of kinase) gene which is frequently overexpressed in cancer and acts as an oncogenic driver promoting aggressive tumor behaviors like metastasis, tissue invasion through increased mobility, and rapid cancer cell growth.

Progression of tumor cell invasion into target distal tissues as metastasis requires a critical early step in the activation of epithelial to mesenchymal transition (EMT), a regulatory program intimately linked to continuous inflammation [48]. EMT allows dissociation of cells from the primary tumor and acquisition of mobility. In a mouse model of pancreatic cancer, EMT was identified as an early event in malignant cell dissemination and the anti-inflammatory dexamethasone suppressed dissemination [62]. The Zeb family of transcription factors (zinc finger E-box binding homeobox) are involved in the regulation of EMT [63]. Zeb1 is a crucial EMT inducer and repressor of E-cadherin, providing stem cell-like features to tumor cells and causing therapy resistance [50]. Its abnormal expression has been reported in pancreatic, lung, liver, colon and breast cancer [48, 50]. Zeb1 is shown here to be greatly inhibited by vitD in mammary tumor cells *in vivo* and *in vitro*. We have previously shown vitD to decrease Zeb1 expression in mammary tumor cells as well as to inhibit chemokine CXCL12 and receptor CXCR4 interaction in lungs as the target organ [33]. The normal CXCL12/CXCR4 signaling is highjacked by cancer cells to drive cancer progression and metastasis; we show here a similar inhibition by vitD of the expression of tumor cell attraction factor CXCL12 in the bone environment, illustrating a similar mode of anti-invasion action for vitD in different target organs.

Another factor impacted by vitD levels in the bone invasion system is nestin (neuroepithelial stem cell protein), a filament protein found in a broad variety of cells such as proliferating vascular endothelial cells, basal cells of the mammary glands, and liver stem cells [36–38]. It is also highly-expressed in many high-metastatic cancers including breast, and is associated with aggressiveness and poor prognosis. Correlation between nestin protein expression and tumor aggressiveness, clinicopathological features, and poor survival rates has been confirmed for many tumors. Nestin is considered a biomarker of invasive phenotype, and is associated to infiltration in glioblastoma, angiogenesis in numerous malignancies and spreading of non-epithelial and epithelial tumors. Overexpression of nestin is linked to aggressive cancer progression and poor prognosis due to its inhibition of cell membrane rigidity that allows extravasation of tumor cells and distal tissue invasion [64].

It is now known that chronic inflammation sustains the tumor microenvironment, allowing cancer cells to bypass apoptosis and proliferate. Inflammation also facilitates establishment of secondary tumors and contributes to treatment resistance. Targeting inflammation is therefore a likely strategy for countering invasion and metastasis. As a role for supplementation of vitD in modifying inflammatory disease is becoming defined, it is clear that vitD status is related to the pathogenesis of skeletal invasion by tumor cells through its implication in countering inflammation factors.

## Financial Support

This work was supported by grant number MOP 10839 from the Canadian Institutes of Health Research to RK.

## Author contributions

Designing research studies: RK; conducting experiments: JL and AL; acquiring data: JL and AL; analyzing data: JL, AC, and RK; providing reagents: RK; writing the manuscript: AC and RK.

## Disclosures

No potential conflicts of interest.

## Data Availability

Some or all data generated or analyzed during this study are included in this published article.

## Notes

### Competing Interest Statement

The authors have declared no competing interest.

