## supplemental figure for "Vitamin D counters bone invasion by mammary cancer through inhibition of inflammation and epithelial-to-mesenchymal transition"

**SUPPLEMENTARY MATERIALS;**

**Supplementary figure 1: Product Summary for Proteome Profiler Mouse Cytokine Array Kit, Panel A.** In red: localisation on the array of the 11 cytokines displaying a large difference in expression between vector-treated and 1, 25$(OH)_2$D -treated cells. Cytokines without red localisation show no significant difference in expression due to treatment.

| | | |
|---|---|---|
| **CXCL13/BLC/BCA-1** | **IL-5** | **M-CSF** |
| **C5a** | **IL-6** | **CCL2/JE/MCP-1 D9-D10** |
| **G-CSF** | **IL-7** | **CCL12/MCP-5 D11-D12** |
| **GM-CSF B7- B8** | **IL-10** | **CXCL9/MIG** |
| **CCL1/I-309** | **IL-12 p70** | **CCL3/MIP-1 α D15-D16** |
| **CCL11/Eotaxin** | **IL-13** | **CCL4/MIP-1 β D17-D18** |
| **ICAM-1 B13 – B14** | **IL-16** | **CXCL2/MIP-2 D19-D20** |
| **IFN-gamma** | **IL-17** | **CCL5/RANTES D21-D22** |
| **IL-1 alpha/IL-1F1** | **IL-23** | **CXCL12/SDF-1 D23-D24** |
| **IL-1 beta/IL-1F2** | **IL-27** | **CCL17/TARC** |
| **IL-1ra/IL-1F3 B21-B22** | **CXCL10/IP-10 D1-D2** | **TIMP-1** |
| **IL-2** | **CXCL11/I-TAC** | **TNF-alpha** |
| **IL-3** | **CXCL1/KC** | **TREM-1** |
| **IL-4** | | |
